# Sleep-wake cycle and behavioral disturbance in a mouse model of frustrative nonreward

**DOI:** 10.1101/2025.08.22.671819

**Authors:** Martina Holz, Rocio C. Fernandez, Mariano Rodriguez, Nina D. Possamai, Alexei L. Vyssotski, Mauricio R. Papini, Horacio de la Iglesia, Ruben N. Muzio, Maria I. Sotelo

## Abstract

Daily time-restricted access to a palatable reward triggers anticipatory behavior that can synchronize an animal’s internal clock and adjust its behavioral rhythms. The unexpected violation of this reward expectancy can induce a negative emotion usually referred to as “frustration”. Frustration responses have been characterized in rodents using a reward downshift paradigm. Although an anxiogenic effect of this reward downshift has been described, the associated long-term locomotor and behavioral patterns remain understudied. Furthermore, changes in the sleep preparatory behavior or in sleep per se associated with sudden reward downshift currently remain underexplored. In this study, we combined wireless electrocorticographic/electromyographic (ECoG/EMG) recordings, accelerometer and video recordings in freely behaving mice to determine sleep-wake, locomotor and behavioral patterns associated with a reward downshift paradigm. Our results show an initial increase in locomotor activity and increased latency to rest following reward devaluation. Furthermore, we found that downshifted mice displayed a reduced number of episodes of rapid-eye-movement sleep (REMs) during the light phase, and increased latency to non-REMs and REMs on the first two days post-shift. Additionally, we found decreased locomotor activity and reduced nest quality over the first two days post-shift compared to unshifted animals. Our results suggest that frustration induces a transient anxiogenic effect followed by a depressive-like state. Of note, our study has the strength of using a novel approach for home-cage behavior analysis that combines machine-learning-based tracking, long-term average body-acceleration, ECoG/EMG recordings, and home-cage nest quality assessment.

## Introduction

Animals display behavioral rhythms that are fundamental to organizing their daily routine and maximizing their capacity to fulfill competing needs such as nutrition, social interaction, and sleep. These rhythms are governed by circadian clocks and, while the light-dark cycle is the principal environmental cycle that entrains circadian clocks, other 24-h nonphotic environmental cues can also entrain circadian rhythms. Among these, the daily time-restricted availability of food can entrain a circadian oscillator and shift an animal’s activity to anticipate the time of reward (Mistlberger, 2009). The daily, predictable availability of a palatable reward, such as sweet or fatty food, can also work as a cue that organizes activity patterns (Muñoz-Escobar, 2019). Studies have shown that this craving for the expected reward can elicit strong anticipatory behavior that can persist even after the reward is suppressed (Angeles-Castellanos et al., 2008).

Several studies have shown that the unexpected omission or reduction of a palatable reward can trigger a negative emotional state, usually referred to as ‘frustration’ or frustrative nonreward (FNR; Amsel, 1992; Papini et al., 2024). In rodents, this effect has been studied mostly using the successive negative contrast (SNC) paradigm (Flaherty, 1996; Papini, 2003). In one version of this paradigm, called consummatory SNC (cSNC), rodents receive access to a highly concentrated sucrose solution during several consecutive days at a fixed time of the day and eventually the solution is unexpectedly downshifted in concentration. Such reward downshift leads to an abrupt reduction in consumption and to the emergence of stress-related responses (Papini, Fuchs, & Torres, 2015). For example, rats exposed to a cSNC task exhibit an increase in locomotor activity, rearing, escape behaviors and a decreased approach to the feeder (Jiménez-García et al., 2016; Norris et al., 2009; Pellegrini & Mustaca, 2000); increased release of stress hormones (Kawasaki, & Iwasaki, 1997; Romero et al., 1995;); reduction in pain sensitivity (Mustaca & Papini, 2005), and increased intake of anxiolytic substances (Manzo et al., 2015). Taken together, this evidence suggests that the unexpected reduction in reward magnitude is accompanied by negative emotion (Papini et al., 2015). Neurobiological evidence shows that some brain areas implicated in FNR, including corticolimbic and hypothalamic regions, (e.g., Abler et al., 2005; Fernandez et al., 2024; Ortega et al., 2013; Tseng et al., 2019; Yu et al., 2014) are also involved in sleep-preparatory behavior and sleep regulation (Hong et al., 2023; Tossel et al., 2023).

We reasoned that the anxiety-related behaviors associated with the sudden reduction of the appetitive reward would disrupt sleep-preparatory behavior, overall sleep-wake patterns, and sleep quality. To test these predictions, we used wild-type male mice implanted with electrodes for wireless electrocorticography (ECoG) and electromyography (EMG) recordings, an accelerometer, and monitored through continuous video recordings to determine sleep-wake architecture and locomotor/activity patterns associated with anticipatory behavior and unexpected reward downshift.

Our results show that a frustrating event such as the sudden and unexpected reduction of a reward induces initially an increase in locomotor activity, in the time spent in the rewarded zone, and in the latency to rest, all compatible with an anxiety-like response. This initial response is followed by longer-term changes compatible with a depressive-like state, including a reduction in the number of episodes of rapid-eye-movement sleep (REMs) during daylight, increased latency to non-REM sleep (NREMs) and REMs, persistently reduced locomotor activity, and reduced nest quality.

## Method

### Subjects

We utilized a total of 23 naïve male, C57 mice (8-12 weeks old at the time of surgery), provided by the IBYME animal facility. They were housed in a controlled environment with a 12-h light/dark cycle, at a temperature of 23 ±1 °C, and had ad libitum access to food, water, and nesting material (one cotton ball and blotting paper). Seven mice were used for the sucrose preference test and 16 for the consummatory reward downshift task. The latter were randomly assigned to the downshift condition (n=8) and the unshifted control condition (n=8). All animal protocols were approved by the Institutional Animal Care and Use Committee (IACUC protocol 001/2023 IBYME-CONICET, Argentina).

Mice were anesthetized with a ketamine-xylazine-acepromacine mixture (100, 10, and 2 mg/kg, respectively; i.p.) and administered a drop of 2% lidocaine as a local anesthetic. They were placed into a stereotaxic frame and were surgically fitted with five miniature screws. Four screws, two in each hemisphere, were used as reference electrodes (frontal: AP=1.5 mm, ML=-1.5 mm, Parietal: AP=−3.5 mm, ML=-2.5 mm), and one screw serving as a ground electrode (AP=-5.9 mm, ML=0). Two electrodes were inserted in the trapezius muscles. All electrodes were pre-soldered to a 7-pin connector and secured to the head using C&B Metabond (Parkell, Edgewood, NY) and dental cement. The skin was sutured with surgical stitches. Animals were injected with meloxicam (5 mg/kg) and yohimbine (0.5 mg/kg) to reduce pain induced by the surgery and were allowed to recover in their home cage for 5-7 days.

### Reward downshift procedure

After recovering from surgery, mice (n=16) were connected to dummy boxes of similar shape and weight to the minilogger recording device (Figure 1A and 1B). They were allowed to habituate to the dummy minilogger a week before experiments started. The experimental protocol was repeated 8 times, each one with a downshifted and an unshifted control pair recorded at the same time. The training procedure was based on Mustaca et al. (2000) with slight modifications (Figure 1B and 1C). Briefly, a sucrose solution (16% concentration for downshifted mice and 2% for unshifted mice) was presented in each animal’s home-cage for 1 h by the end of the dark phase [zeitgeber time (ZT) 23-ZT0, with ZT0 the time of lights ON] for 10 consecutive pre-shift sessions. On session 11 (first post-shift session) the 16% sucrose solution was downshifted to 2% sucrose for downshifted animals, to match that of unshifted controls that were always exposed to 2% sucrose. There were 5 post-shift sessions (sessions 11-15) during which both groups had access to 2% sucrose. ECoG-EMG and accelerometer data were recorded on pre-shift sessions 1, 9, and 10, and on post-shift sessions 11, 12, and 15 (Figure 1B). Video recordings were stored for all days starting 10 min before session onset and lasting 23 h.

**Figure 1.**
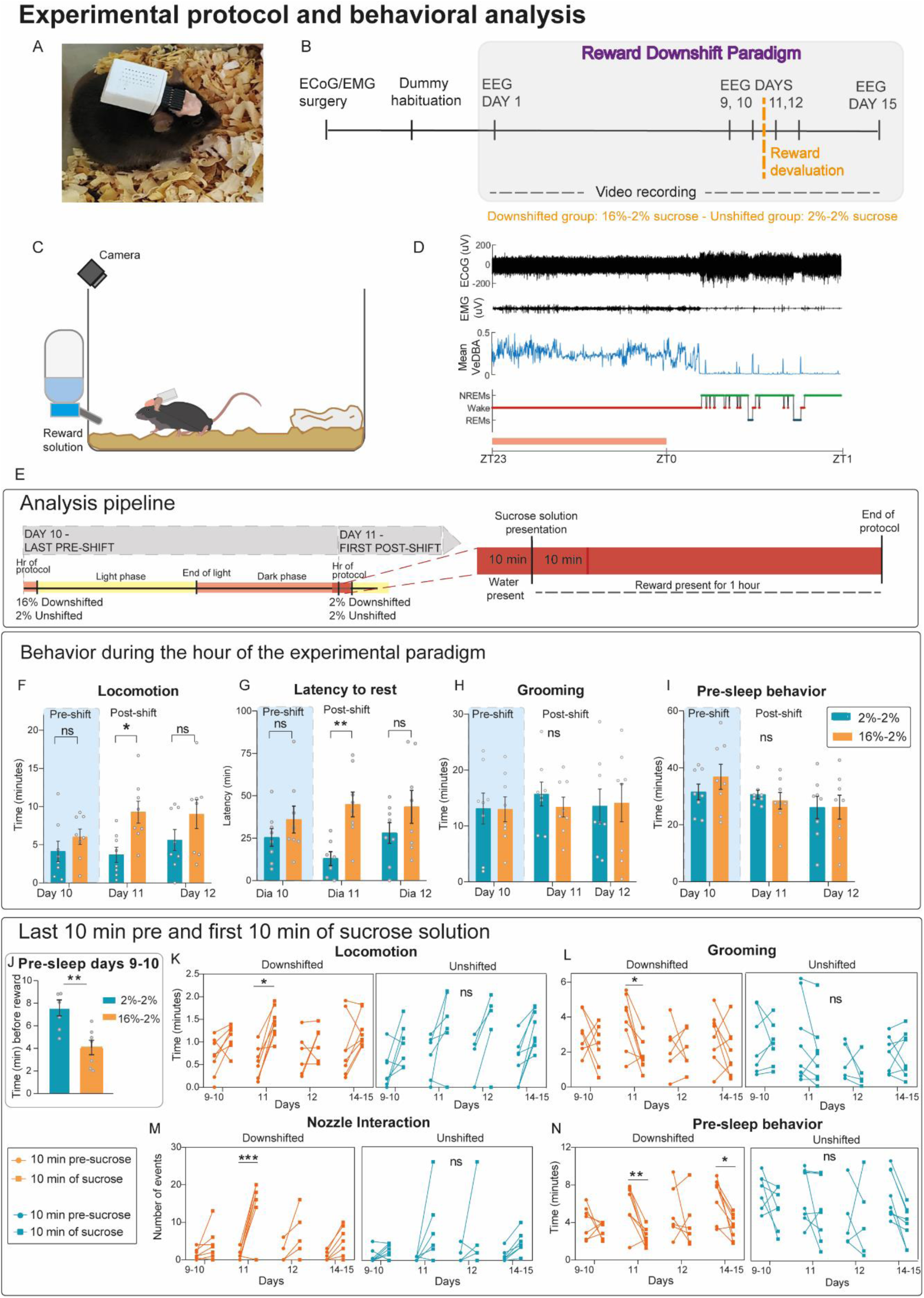
A) Photograph of an ECoG/EMG implanted C57 WT mouse wearing the Neurologger 2A. B) Schematics of the Reward Downshift Paradigm including the previous surgery and dummy habituation. The 16% reward is presented on days 1-10 (pre-shift phase) and is then downshifted to 2% on days 11-15 (post-shift phase) for the ‘downshifted’ group. C) Schematics of the animal home-cage and the mouse wearing the recording device and receiving the reward solution during the protocol. D) ECoG, EMG, mean VeDBA calculated from x,y,z acceleration and hypnogram of a downshifted mouse during the protocol hour (ZT23-ZT0) and the first hour of daylight (ZT0-ZT1). E) Schematics of the analysis pipeline on days 10 and 11 of the experimental paradigm (as example) showing the hour while the sucrose is present and the remaining hours of the light-phase and dark-phase that were recorded (12h and 10h respectively). On the right side see zoomed-in schematics of the 10 min windows analyzed prior to and following sucrose presentation. F-I) manually analyzed behavior during the hour of the paradigm on days 10, 11 and 12. F) Locomotion, n=8 mice, two-way RM ANOVA, p(Treatment) = 0.0424, p(Treatment x Day) = 0.15. G) Latency to rest, n=8 mice, two-way RM ANOVA, p(Treatment) = 0.0018, p(Treatment x Day) = 0.31. H) Grooming, n=8 mice, two-way RM ANOVA, p(Treatment) = 0.8, p(Treatment x Day) = 0.82. I) Pre-sleep behavior, n=8 mice, two-way RM ANOVA, p(Treatment) = 0.51, p(Treatment x Day) = 0.73. J) Last 10 min prior to sucrose presentation on days 9-10 showing the difference in ‘pre-sleep’ behavior in the downshifted and unshifted animals. n=8 mice, unpaired t-test, p = 0.0033. K-N) Behavior in downshifted (left) and unshifted animals (right) in the 10 min before sucrose presentation compared to the first 10 min with the sucrose present in the cage. K) Locomotion, n=8 mice, three-way RM ANOVA, p(Treatment) <0.0001, p(Treatment x Day) = 0.66, p(Treatment x 10min-window) = 0.56, p(Treatment x x Day x 10min-window) = 0.22. L) Grooming n=8 mice, three-way RM ANOVA, p(Treatment) = 0.0085, p(Treatment x Day) = 0.25, p(Treatment x 10min-window) = 0.23, p(Treatment x x Day x 10min-window) = 0.74. M) Nozzle Interaction, n=8 mice, three-way RM ANOVA, p(Treatment) = 0.0003, p(Treatment x Day) = 0.0007, p(Treatment x 10min-window) = 0.2, p(Treatment x x Day x 10min-window) = 0.11. N) Pre-sleep, n=8 mice, three-way RM ANOVA, p(Treatment) = 0.0002, p(Treatment x Day) = 0.21, p(Treatment x 10min-window) = 0.72, p(Treatment x x Day x 10min-window) = 0.35. ns, p > 0.05; ∗, p < 0.05; ∗∗, p <0.01; ∗∗∗, p <0.001.

For each mouse, fresh sucrose solution was prepared each day on a w/w basis by diluting 16 g (or 2 g) of sucrose for every 84 g (or 98 g) of water. One hour before the start of each session, food pellets and the water bottle were removed from the cage and a bottle with the sucrose solution replaced the water bottle (Figure 1E). If ECoG/EMG was to be recorded on that day, mice were disconnected from their dummy loggers and connected to the miniloggers (Neurologger 2A, Evolocus, Tarrytown, NY) around 30-20 min before inserting the sucrose bottle. The miniloggers were set to record ECoG/EMG and accelerometer data (x, y, z axis) at 400 Hz for 23 h (Figure 1D, example). Web cameras (1080P, Angetube, Shenzhen, China) located on top of each cage were started at least 10 min before changing the solution. Videos were recorded using iSpy software (https://iSpyConnect.com) at 15 fps. To produce higher quality videos for example purposes, we included an additional IR camera (2MP, ArduCam, Kuala Lumpur, Malaysia) on the opposite side of the cage phasing the reward solution for 2 unshifted and 2 downshifted animals (see Video S1). We used Lightworks 2022.3 free software (http://lwks.com/) to produce a high-speed (80x) video recorded only while the solution was present.

Reward solutions were offered for 1hr (ZT23 to ZT0), and they were left in the cage until the lights of the room were turned on for the start of the light phase. When lights turned on at ZT0, the reward solutions were removed from each cage, they were weighed, and the water bottle and food pellets were returned to the home cage. Web-cameras and miniloggers were set to keep recording for the rest of the day (22 h on) and videos were saved the next day at around ZT22 and before starting a new day of training.

### Sucrose preference test

Mice (n=7) were exposed to two bottles with different concentrations of sucrose in their home cage for 24 h and 5 days a week (Figure S1A). At the beginning of the day (ZT0) each bottle of solution was removed from the cage, weighed, and replaced by fresh solution for another 24 h. During weekends, animals were allowed to hydrate with tap water for two days before the test continued. The test lasted for three weeks and the combination of sucrose solution was as follows for each animal: week 1: 32% and 4% sucrose, week 2: 16% and 2% sucrose, week 3: 2% sucrose and 0% (deionized water). Exposure to 32%-4% and 16%-2% sucrose was counterbalanced across animals, with half the mice receiving 16%-2% on week 1 and the rest on week 2. We did not test preference between 4% and 0% concentration since preliminary results indicated a stable preference for the 16%-2% combination.

### Polysomnographic analysis

We analyzed ECoG-EMG signals collected at 400 Hz during sessions 10, 11, 12, and 15 of the experimental protocol using the free AccuSleep interface for MATLAB (Figure 3 and S2B-I). Recordings were first automatically scored based on sample scoring by a human observer and the automatic analysis was manually corrected where needed. Recordings were organized in 5-s epochs into wakefulness, NREMs, and REMs. Wakefulness was defined as desynchronized, low-amplitude ECoG signal occurring concomitantly with high tonic EMG signal with phasic bursts. NREMs periods were characterized as synchronized, high-amplitude, low-frequency (0.5–4 Hz) ECoG signals with reduced EMG activity compared to wakefulness. REMs periods were defined as ECoG signals with a dominant theta band (6–9 Hz) with very low EMG activity. The extracted hypnograms for each animal and day were processed using custom MATLAB software to extract the percentage, the number of episodes, and the mean episode duration for each state. Recordings were obtained 1, 2, and 12 h inside the light phase period, and 10 h inside the dark phase period (recordings were about 23-h long for each day and the first hour was recorded during the reward downshift protocol). Power spectrograms were calculated for each state using a Fast Fourier Transformation and extracting band signals for delta (0.5-4 Hz), theta (5-9 Hz), sigma (11-15 Hz), beta (15-30 Hz), and gamma (30-100 Hz) power. To extract power, we first removed any existing movement artifacts from the ECoG/EMG signals by manual inspection with the AccuSleep interface and custom MATLAB software. Then, relative values were extracted by dividing by the total power of wakefulness during the 10h of the dark phase for each recording of each animal. We excluded two animals from spectrogram analysis, one from the downshifted and one from the unshifted condition, due to noisy ECoG signals containing movement artifacts.

**Figure 2.**
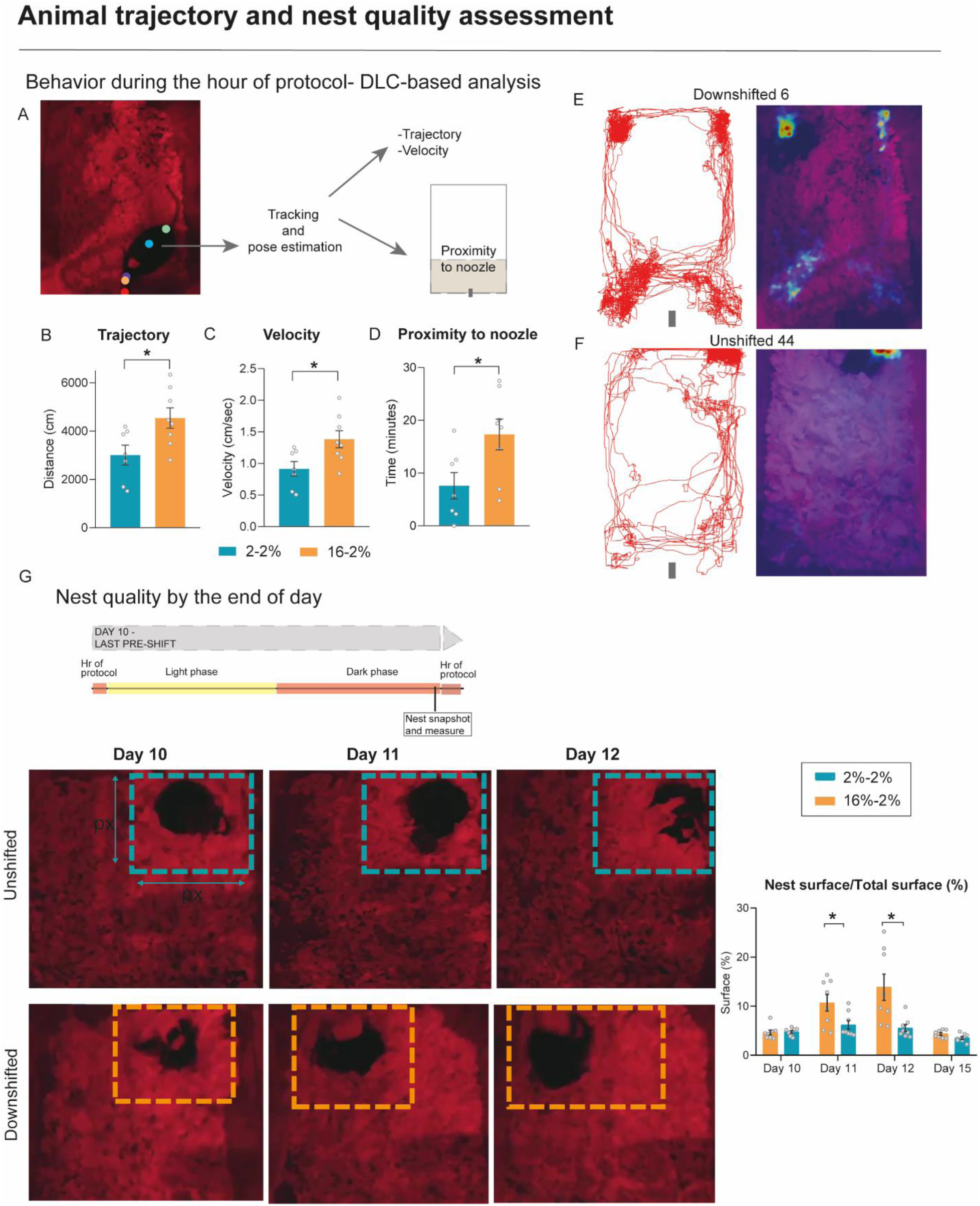
A) Schematics of the DLC-tracking and key points used. The light-blue point on the middle of the back was chosen to track the animal and extract trajectory, velocity and proximity to the reward nozzle. B-D) Trajectory, velocity and time spent in the rewarded zone during day 11 for downshifted and unshifted animals during the hour of the experimental paradigm. B) Trajectory, n=8 mice, unpaired t-test, p = 0.0228. C) Velocity, n=8 mice, unpaired t-test, p = 0.0227. D) Proximity to nozzle, n=8 mice, unpaired t-test, p = 0.0267. E) Example trajectory (left) and heat map (right) of a downshifted and F) unshifted animals on day 11. G) Schematics of day 10 (as example) showing the moment where the snapshot was chosen to evaluate nest quality (prior to reward presentation of the following day). Below, see example snapshots used for days 10-12 for an unshifted (top) and a downshifted animal (down). On the right side, see results of the % surface occupied by the nest in the home-cage used as an indirect measure of nest quality. % nest surface/ total surface, n=8 mice, two-way RM ANOVA, p(Treatment) = 0.0085, p(Treatment x Day) = 0.0011. ns, p > 0.05; ∗, p < 0.05; ∗∗, p <0.01.

**Figure 3.**
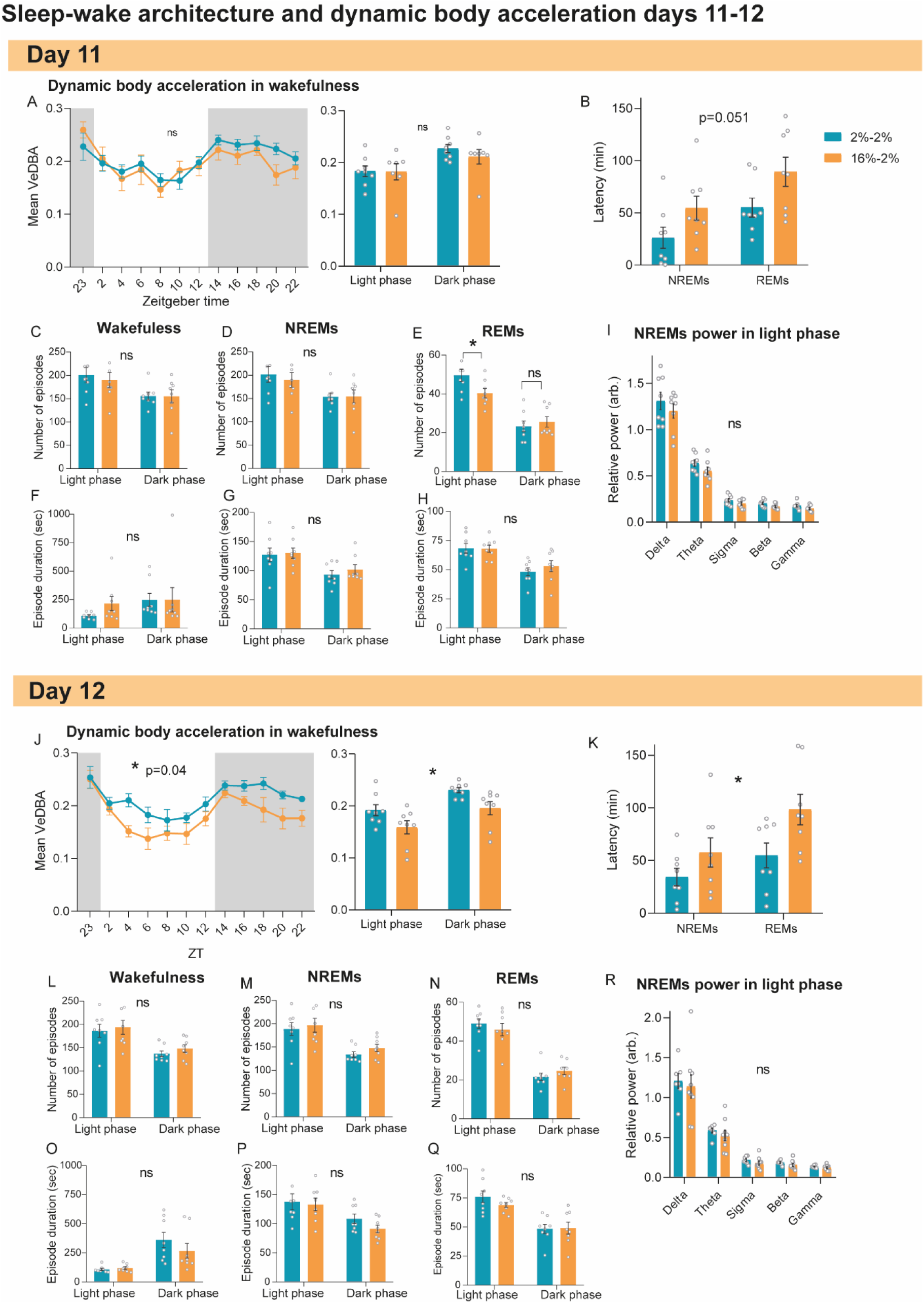
A) Dynamic body acceleration or VeDBA calculated for the hour of protocol and 2 hour means (left) on day 11 (first day of down-shift for downshifted animals), n=8 mice, two-way RM ANOVA, p(Treatment) = 0.69, p(Treatment x Hour) = 0.63. On the right side, see mean values in the 12h of light phase and 10 h of dark phase post-reward devaluation, n=8 mice, two-way RM ANOVA, p(Treatment) = 0.61, p(Treatment x Phase) = 0.098. B) Latency to NREMs and REMs sleep from ZT0 and after the reward is removed from the cage on day 11, n=8 mice, two-way RM ANOVA, p(Treatment) = 0.051, p(Treatment x Stage) = 0.66. C-E) Number of episodes of wakefulness, NREMs and REMs during the 12h of light phase and 10h of dark phase following reward devaluation. C) Wakefulness, n=8 mice, two-way RM ANOVA, p(Treatment) = 0.75, p(Treatment x Phase) = 0.61. D) NREMs, n=8 mice, two-way RM ANOVA, p(Treatment) = 0.74, p(Treatment x Phase) = 0.54. E) REMs, n=8 mice, two-way RM ANOVA, p(Treatment) = 0.35, p(Treatment x Phase) = 0.0022. F) Mean episode duration of wakefulness, NREMs and REMs during the 12h of light phase and 10h of dark phase following reward devaluation. F) Wakefulness, n=8 mice, two-way RM ANOVA, p(Treatment) = 0.52, p(Treatment x Phase) = 0.8. G) NREMs, n=8 mice, two-way RM ANOVA, p(Treatment) = 0.45, p(Treatment x Phase) = 0.71, H) REMs, n=8 mice, two-way RM ANOVA, p(Treatment) = 0.98, p(Treatment x Phase) = 0.15. I) Mean relative power in absolute values for delta-gamma bands in NREMs during the light phase, n=8 mice, two-way RM ANOVA, p(Treatment) = 0.19, p(Treatment x Frequency) = 0.72. No significant differences were found although a strong tendency to increased beta power in the downshifted animals was present. J-R) See same results as previously described for day 12. J) VeDBA calculated for the hour of protocol and 2 hour means (left) on day 12, n=8 mice, two-way RM ANOVA, p(Treatment) = 0.48, p(Treatment x Hour) = 0.0405. On the right side, see mean values in the 12h of light phase and 10 h of dark phase post-reward devaluation, n=8 mice, two-way RM ANOVA, p(Treatment) = 0.86, p(Treatment x Phase) = 0.0338. K) Latency to NREMs and REMs sleep from ZT0 and after the reward is removed from the cage on day 12, n=8 mice, two-way RM ANOVA, p(Treatment) = 0.0429, p(Treatment x stage) = 0.27. K-N) Number of episodes of wakefulness, NREMs and REMs during the 12h of light phase and 10h of dark phase on day 12. K) Wakefulness, n=8 mice, two-way RM ANOVA, p(Treatment) = 0.54, p(Treatment x Phase) = 0.22. L) NREMs, n=8 mice, two-way RM ANOVA, p(Treatment) = 0.59, p(Treatment x Phase) = 0.66. M) REMs, n=8 mice, two-way RM ANOVA, p(Treatment) = 0.58, p(Treatment x Phase) = 0.54. N-P) Mean episode duration of wakefulness, NREMs and REMs during the 12h of light phase and 10h of dark phase on day 12. N) Wakefulness, n=8 mice, two-way RM ANOVA, p(Treatment) = 0.39, p(Treatment x Phase) = 0.28. O) NREMs, n=8 mice, two-way RM ANOVA, p(Treatment) = 0.41, p(Treatment x Phase) = 0.31. P) REMs, n=8 mice, two-way RM ANOVA, p(Treatment) = 0.49, p(Treatment x Phase) = 0.35. I) Mean relative power in absolute values for delta-gamma bands in NREMs during the light phase, n=8 mice, two-way RM ANOVA, p(Treatment) = 0.52, p(Treatment x Frequency) = 0.98. ns, p > 0.05; ∗, p < 0.05.

### VeDBA calculation

To estimate general locomotor activity, we calculated a dynamic body acceleration (VeDBA) parameter combining 3-axial dynamic acceleration (Shepard et al., 2008; Wilson et al., 2019). Briefly, we employed custom MATLAB software to analyze 3-axial acceleration recorded by the miniloggers at 400 Hz. We smoothed the raw acceleration data to remove static acceleration by subtracting a running mean of 2 s for each recorded axis (x, y, z). Then we employed the VeDBA equation (Wilson et al., 2019) to compute dynamic acceleration resulting from body movement using absolute values, and we calculated mean values in 1, 2, and 12 h inside the light period, and 10 h inside the dark period on days 10, 11, 12, and 15 (Figure 3A, J and S2B, F).

### Video analyses

#### BORIS ethogram

We centered on manual analysis of the hour of the protocol where the reward solution was present for downshifted and unshifted mice and the 10 min window before the solution was changed (before ZT23). This rendered a total of 1 h and 10 min of analysis (Figure 1E).

The analysis was divided as follows: (1) a 1-h analysis while the solution was present (ZT23-ZT0) using the ethogram from Table 1 (modified from Sotelo et al., 2022; n=8 mice for unshifted and downshifted mice) (Figures 1F-I and S1C). (2) an analysis of the 10 min before changing the sucrose solution (while food and water were present) and the first 10 min after the change (sucrose solution present, food and water absent) with a modified ethogram that included ‘Nozzle Interaction’ (Table 1, extended) (Figures 1J-N and S1D-I). The idea behind this second analysis was to characterize predicted behavioral shifts associated with reward anticipation and to compare them with behavior after the unexpected reward downshift. This analysis was done utilizing BORIS software (Friard & Gamba, 2016) and a detailed behavioral annotation by manual observers in 1-s periods. Observers were blind to the assignment of animals to the conditions. One animal was excluded from the unshifted group and one from the experimental group since their pre-sucrose video recordings were shorter than 10min (n=7 unshifted and n=7 downshifted mice).

**Table 1.** Ethogram used to classify behavior during the hour of the protocol. In grey, see extended ethogram used for the 10 min prior to reward and first 10 min of reward behavioral analysis including ‘Nozzle Interaction’.

| Behavior | Description |
| --- | --- |
| Drinking | The mouse is licking the bottle spout. The tongue could be seen but it lasts less than one second. |
| Resting | The mouse is still and inactive, generally in the nest and probably sleeping. It usually adopts a characteristically enrolled position or with its face hidden in some part of its body. It can include brief twitches (< 4 s long). |
| Locomotion | The mouse displaces inside the cage, walking or running. It could be fast or slow. |
| Pre-sleep | The mouse is engaged in nest-building (manipulating nesting material, digging, fluffing or "sniffing" the material) or it is staying 'still and alert' inside the nest (not resting, visibly awake). The mouse can also engage in grooming behavior (only grooming in the nest is included). |
| Rearing | The mouse stands in its hindlegs and in equilibrium or supports with its front legs pressed to the cage. It could include a small jump. |
| Grooming | The mouse is cleaning its body with its tongue or scratching with its hindlegs. |
| Nozzle Interaction | Animal is biting or sniffing the snout but not drinking. |
| Other | Any other behavior that is not encapsuled by the previously mentioned or it is not of interest. |

#### Deep-Lab-Cut based automated analysis

The 1-h videos recorded while the sucrose solution was present were also analyzed with an automated machine-learning-based approach using a DeepLabCut (DLC, version 2.3.11) single animal model (Mathis et al., 2018). Briefly, we trained a DLC model to detect singly housed mice in their home cage during the light and dark phases using our created video recordings from the experimental protocol plus other videos from 24-h recordings in the home-cage. We randomly extracted 20 frames from 27 different videos. On each of the frames (480x640 pixels), we manually annotated the tip and the end of the sipper tube in the cage, and the nose, a point in the middle of the back, and a point at the end of the back (near the base of the tail) on the mouse. We used a standard 95% of the frames for training. We used a ResNet-50-based neural network with default parameters and we trained the model for 580,000 iterations. Then we evaluated the model using DLC software publicly available on Google colab (https://colab.google/) notebooks (train error: 2.95 pixels, test error: 5.6 pixels, p-cutoff = 0.4) and we analyzed and filtered the video data from sessions 10, 11, and 12 of the reward downshift protocol from ZT23 to ZT0. We extracted x and y coordinates for each part of the body analyzed and we specifically used the middle back point to detect and track the animal, since this was the most stable point in all cases. We only used data points from frames with more than 0.7 likelihood to detect trajectory paths. Those below 0.7 were replaced by the previous coordinate with a likelihood greater than 0.7. We used custom MATLAB and Python software to extract heatmaps, trajectory maps, mean velocity and time spent close to the bottle nozzle (frames where the animal was detected within 1/3 of the cage that was closer to the reward) during the hour of the protocol on sessions 10, 11, and 12 (Figure 2A-F and S1J-Q). We had to exclude an unshifted animal from the analysis that had built its nest next to the sipper tube because this would significantly affect the time spent close to the bottle. We also excluded sessions 10 and 12 for another downshifted animal due to low quality of tracking that probably corresponded to very low illumination in the cage.

### Nest quality assessment

We quantified in-home nest quality in the home-cage by approximating a rectangle that encapsulated the 2D surface of the nest (Figure 2G and S2J). Briefly, we used a snapshot of the cage taken from the video-recording during ZT22 on days 10, 11, 12, and 15 picking a moment where the animal was sleeping or staying still in the nest. We store the snapshots for each mouse and two independent observers used Photoshop CS2 Portable (Adobe, 2005) to draw a rectangle surrounding the limits of the nest, extracting the length and width measurements in pixels. The surface of the nest was then calculated by relating the previous measurements to the DPI (dots per inch) of each image. To express this information, the total surface area of the image was related to that of the nest, to establish the "% surface of the nest”.

### Individual variability calculation

Individual differences were analyzed to explore variability in responses following reward devaluation. Analyses were conducted using R (R Core Team, version 4.4.1). Missing data from one animal on day 10 were imputed using the group mean (mouse 43 for “Trajectory,” “Velocity,” and “Proximity to nozzle”). Another animal was excluded from analysis due to missing data on day 11. Individual differences for each variable were quantified by calculating log-ratios, log(day11/day10), where 0 indicates no change in response, positive values indicate an increase (red), and negative values indicate a decrease (blue). Supplementary Figure S2A illustrates the variability observed across individuals for multiple variables following reward devaluation.

### Statistical analysis

We used GraphPad Prism software (version 8.0.1) to analyze and plot behavioral, ECoG/EMG and accelerometer data. To assess group differences in behavioral variables and solution intake during the h of reward solution presentation across days, a two-way repeated-measures (RM) ANOVA (Treatment x Day) was performed (Figure 1F-I and S1B, C). Post hoc multiple comparisons were carried out using Fisher’s LSD test. We also used two-way RM ANOVA (Treatment x Phase) to compare number and duration of episodes of wakefulness, NREMs and REMs and VeDBA during wakefulness across different days pre-and post-shift (Figure 3A-H, 3J-Q). We used two-way RM ANOVA (Treatment x Frequency) to test for differences in spectrogram frequency bands on different days (Figure 3I, R).

To examine anticipatory and immediate effects related to sucrose solution shift, behavioral variables were analyzed during the 10 min before and following reward presentation. We used three-way RM ANOVA with the following factors: days (9-10, 11, 12, 14-15), 10 min-window (10 min pre-sucrose, 10 min of sucrose) and treatment (downshifted or unshifted), followed by post-hoc multiple comparison Tukey tests (Figure 1K-N and S1H-I). We separately used unpaired t-tests only on day 9-10 (averaged values) and day 14-15 to assess possible differences related to fix-time palatable reward presentation centering on Locomotion, Rearing and Pre-sleep behavior between downshifted and unshifted animals during the 10 min pre-sucrose (Figure 1J and S1D-G).

We used a one-way RM ANOVA for the sucrose preference test with Tukey post-hoc tests (Figure S1A) and a two-way RM ANOVA for the nest quality assessment across days (Figure 2G, Treatment x Day).

We used unpaired t-tests to examine trajectory, velocity and proximity to nozzle on different days (Figure 2B-D and S1J-L, S1N-P).

The Geisser-Greenhouse correction was applied when necessary, and residual normality was assessed using QQ plots. All statistical tests were conducted using an alpha level of 0.05 but with correction for multiple comparisons.

We used R statistical software to compare and plot individual differences between days 11-10 in the downshifted mice (Figure S2A).

## Results

### Unexpected reward downshift induces persistent behavioral and locomotor activity changes in mice during post-shift phase

To determine the effects of frustration on daily behavioral patterns and locomotion, downshifted mice received a high reward consisting of a 16% sucrose concentration during ten consecutive days, which was suddenly and unexpectedly changed to a lower concentration (2%) on day 11 and until the end of the protocol (day 15); unshifted mice received 2% sucrose throughout the protocol. Consistent with previous literature, we found that both downshifted and unshifted animals displayed increased drinking behavior immediately following reward presentation in the pre-shift phase (Figure S1H).

On the first day of downshift (day 11), we detected a reduction in solution intake following reward devaluation that did not reproduce a contrast effect (Figure S1B). However, we did detect significant changes in locomotor variables and time spent in the rewarded zone by the downshifted animals, which are changes that have been previously associated with emotional frustration (see Discussion). Downshifted mice (16%-2%) showed increased latency to rest in their nests and increased locomotor activity during the hour of the protocol compared to unshifted mice that experienced no change (2%-2% sucrose) (Figure 1F-G, Video S1).

Moreover, machine-learning based tracking showed that downshifted mice had a significant increase in their covered trajectory and mean velocity during the first day of downshift while the reward was present, and they spent an increased amount of time in the rewarded zone (Figure 2A-F). These differences were not present on the last day pre-shift nor on the second day post-shift (Figure 1F-G, Figure S1J-P), suggesting that the sudden reduction of the reward induced the changes. We did not detect differences in pre-sleep behavior, rearing or grooming behavior during the hour of the protocol on any of the analyzed days (Figure 1H-I, Figure S1C,D).

To determine anticipatory behavior during pre-shift and post-shift of the reward downshift paradigm, we used a similar modified version of the ethogram (Table 1, extended) and compared a 10 min window before the solution was changed and the first 10 min with the solution present in the home-cage (Figure 1E). On the last days of the pre-shift phase, downshifted animals receiving 16% sucrose, showed significantly decreased time spent on ‘pre-sleep’ behavior in the 10 min before receiving the reward solution compared to unshifted animals receiving 2% sucrose (Figure 1J) and a tendency to spend more time in active ‘locomotion’ (Figure S1F). The difference in pre-sleep behavior between downshifted and unshifted mice was not present after downshift on days 14-15 (Figure S1E, G), which suggests an existing pre-shift difference in anticipatory behavior between animals receiving a highly palatable reward (16% sucrose) and those receiving a more diluted reward (2% sucrose). After downshifting, we found an immediate decrease in the time spent on grooming and pre-sleep behaviors and a strong increase in the number of interactions with the bottle nozzle (‘nozzle interaction’) and in ‘locomotor’ behavior during the first 10 min post-downshift while the devalued solution was present, compared to the 10 min pre-reward (Figure 1K-N). We also found a reduction in total time spent on ‘pre-sleep’ behaviors on day 11 on the first 10 min post-downshift that was still apparent on days 14-15 (Figure 1N). These results suggest increased anxiety-related behavior associated with reward downshift and possibly increased time spent in attentive arousal and reduced time spent preparing for sleep.

### Frustration induced by reward devaluation induces sleep-wake disturbances and it is associated with reduced body acceleration and poor nest quality in mice

We determined sleep-wake architecture in male mice (n=8 unshifted 2-2%, n=8 downshifted 16-2%) during specific days pre-shift (10) and post-shift (11,12,15). Downshifted mice displayed a reduced number of episodes of REM sleep during the light phase of the first day post-shift (day 11) with no changes in the number of episodes of wakefulness or NREMs nor on episode duration during any stage (Figure 3C-H). Of note, there was a tendency for most downshifted animals to display shorter episodes of wakefulness during the dark phase that rendered no significant differences due to subject variability (Figure 3F and S2A). We found no differences in the total amounts of wakefulness or sleep, although a tendency towards more wakefulness and less REMs during the light phase was present (data not shown). Downshifted mice also showed a strong tendency to have increased latency to NREMs and REMs on the first day post-shift (Figure 3B). No differences were found in mean body acceleration throughout day 11 (Figure 3A).

On the second day post-shift (day 12 of the protocol), downshifted mice showed a significant increase in their latency to NREM and REM sleep (Figure 3K) compared to control unshifted mice. No differences were found in the number of episodes or episode duration for this day (Figure 3L-Q) nor in total amounts of wakefulness or sleep (data not shown). Notably, we found an overall decrease in the mean dynamic body acceleration (mean VeDBA) across day and night (Figure 3J) measured with the accelerometer of the minilogger device.

In general, we found no significant differences in animal spectrograms throughout the pre-shift and post-shift days analyzed for the paradigm (Figure 3I, R), although of note, there was strong subject variability and we found a strong tendency to reduced beta power during NREMs in the light phase on the first day of downshift (ns, p=0.059, Figure 3I and S2A).

We did not observe significant differences in sleep-wake architecture or mean body acceleration on day 10 pre-shift and day 15 post-shift which suggests that the observed differences on days 11 and 12 were strictly related to reward devaluation (Figure S2B-I).

Next, we analyzed nest quality in the home-cage specifically at ZT22 (the hour before the protocol started) by semi-randomly choosing a snapshot while each mouse was lying still or asleep in the nest. We measured the surface of the nest (pixels in the x-y axis) as an indirect measure of nest structure, as we preliminary observed that downshifted mice were presumably showing nests of a less consolidated nature. As we predicted from preliminary observation, downshifted mice were sleeping in lower-quality built nests with scattered materials by the end of the first-and second-days post-shift (days 11 and 12 of protocol, Figure 2G). This was evidenced by a larger 2D surface compared to unshifted mice on the same days, and also compared with the same downshifted mice on day 10 (last day pre-shift). We found no differences in nest surface on day 15 between downshifted and unshifted mice (Figure 2G and S2J).

## Discussion

In this study we used a sucrose reward downshift paradigm resembling consummatory SNC that had been previously characterized in rodents, to assess sleep-wake and long-term behavioral and locomotor parameters associated with frustrative nonreward (FNR). Consistent with our prediction, prior to downshift we found decreased time spent in pre-sleep behavior, including nest-building, grooming and staying still and alert in the nest. We also found a tendency to spend increased time in locomotor activity consistent with anticipatory behavior in a time window associated with reward presentation. The lack of a stronger difference in anticipatory locomotor activity between downshifted and unshifted animals could be due to the fact that although unshifted animals were receiving a less concentrated sucrose solution, they were still receiving a reward and therefore exhibiting anticipatory behavior. Previous experimental procedures had mostly compared between a fixed-time presentation of the reward and either a random time presentation or a control receiving no reward (Mistlberger, 2009; Muñoz-Escobar, 2019). All in all, our results support previous findings regarding the salience of palatable food as a circadian clock modulator. On the other hand, and also consistent with our hypothesis and previous literature, we found increased locomotor parameters and trajectory associated with reward downshift that are suggestive of increased anxiety. This increased locomotor response is also apparent on the first 10 min window following the solution presentation along with increased interaction with the bottle nozzle, such as snout biting or sniffing. This result suggests that animals take a first time window of increased attention and evaluation of the reward and possibly increase their ‘search’ behavior. Previous literature had also found increased locomotion in rodents including mice and increased distance travelled following reward downshift in open-field under controlled test settings and short-time windows (10-20 min, Fernandez et al., 2024; Naik et al., 2024; Pellegrini & Mustaca, 2000). In contrast, Pellegrini and Mustaca (2000) had found a reduced approach to the feeder in rats following reward devaluation in a consummatory SNC task with solid food. Our mice spent increased time in the rewarded zone, which suggests that contrary to rats, mice do not experience an avoidance reaction but rather invigorate their search response.

To study sleep-wake and long-term locomotor parameters associated with FNR, we combined this study with ECoG/EMG and accelerometer recordings obtained with wireless minilogger devices. Notably, we found increased latency to rest and delayed NREMs and REMs onset as well as a reduction in the number of REMs episodes during the light phase following reward downshift and compared to unshifted animals. The finding that frustration induced delayed sleep onset appears consistent with some of the past literature on negative emotion impacts on sleep (Krizan et al., 2023). Regarding the changes found in REMs, changes associated with REMs including the number of episodes and REMs latency have been found associated with increased stress levels and corticosterone concentration or with mental pathology (Baglioni et al., 2016; Nollet et al., 2019; Yasugaki et al., 2019; 2025). In most cases the number of REMs episodes has been found to be increased under stress, however, most previous studies have been based on chronic stressors and unpleasant stimulus presentation rather than a palatable reward devaluation or omission. Interestingly, we also found decreased levels of body acceleration throughout the second day and night post-shift. The increased latency to NREMs and REMs and decreased acceleration suggests a long-term depressive-like state associated with the reward downshift. Finally, the possible depressive-like state of downshifted animals is further supported by the analysis of nest quality that revealed an increase in nest surface consistent with more scattered nesting material or a less-consolidated nest structure over the first-and second-nights post-shift. Notably, a recent experiment conducted in mice by Naik et al. (2024) under another FNR paradigm called Alternate Poking Reward Omission (APRO) failed to detect depressive-like behavior following reward devaluation using forced swimming test or sucrose preference test. The differences could be related to multiple factors that were dissimilar to our experimental protocol. Namely, the investigators used water as reward in thirsty animals while in our case we used a palatable reward in animals that were not food or liquid deprived. Also, the team tested the animals in separate arenas, not in the home-cage and the paradigm involved a random omission of the reward in two consecutive days and in a progressive manner (first day they omitted reward 50% and second day 80% of the time). Since we found the reduced acceleration effect mostly post-protocol and in a large period of time in the home-cage, it cannot be discarded that the APRO protocol could still be inducing this long-term effect. Furthermore, our results from in-home nest quality assessment and mean dynamic body acceleration levels support previous findings that suggest that these effects are long-lasting and persistent even when the stressor is removed (Nollet et al., 2019). Interestingly, we did not find differences in locomotor parameters such as trajectory and velocity of the downshifted animals on the second day of downshift while the reward was present, suggesting that the initial hyperkinetic response that we found on the first day had then declined and had been superseded by a lethargic state.

### Final remarks

The present study tried to determine short-term and long-term natural behavior and sleep-wake parameters around a reward downshift paradigm that induces an emotional response in mice similar to human frustration. Our results constitute a first fundamental step that assess long-term effects in a well-characterized and controlled protocol of frustrative nonreward. Moreover, we have established a pioneering approach for home-cage and long-term behavioral assessment, employing wireless technology, machine-learning based tracking and in-home nest quality rating. Future work should focus on the cerebral substrates that underlie this emotional reaction and aim towards the understanding of the link between frustration and sleep and circadian regulatory mechanisms.

## Data availability

Data sets used will be publicly available upon publication. Any additional queries should be directed to the lead author, Ines Sotelo

## Author contributions

M.H., and M.I.S. conceived and designed the study. M.H., N.D.P., and M.I.S. performed research. M.H., N.D.P., R.C.F., M.R., and M.I.S. analyzed data. A.L.V. contributed hardware. M.H., R.C.F., M.R., H.I., M.P., R.N.M., and M.I.S. wrote the manuscript, with feedback from all authors. M.I.S., R.N.M. and H.I. supervised the study.

## Acknowledgments

We would like to thank Agustina Gomez Laich for her help and guidance with VeDBA calculation. Diego Gelman and Martina Belmonte for kindly heling with mice breeding and donation. Ernesto Gulin and Natalia Vasta from animal facility services for helping with the anesthetic protocol, general mice well-being and for providing us with materials for our animal home-cages. Special thanks to Horacio de la Iglesia for all his mentoring and patience during this process.

---

**Video S1**. Example fast-played recording of a downshifted animal showing the behavior right before the water and food are removed and the reward solution is placed on day 10 (16% sucrose solution) and on day 11 (2% solution for the first time) for the same downshifted mouse. Speed is 80x real speed and video was recorded with an IR camera located on top of the nest and opposite to the reward spout.

**Figure S1.**
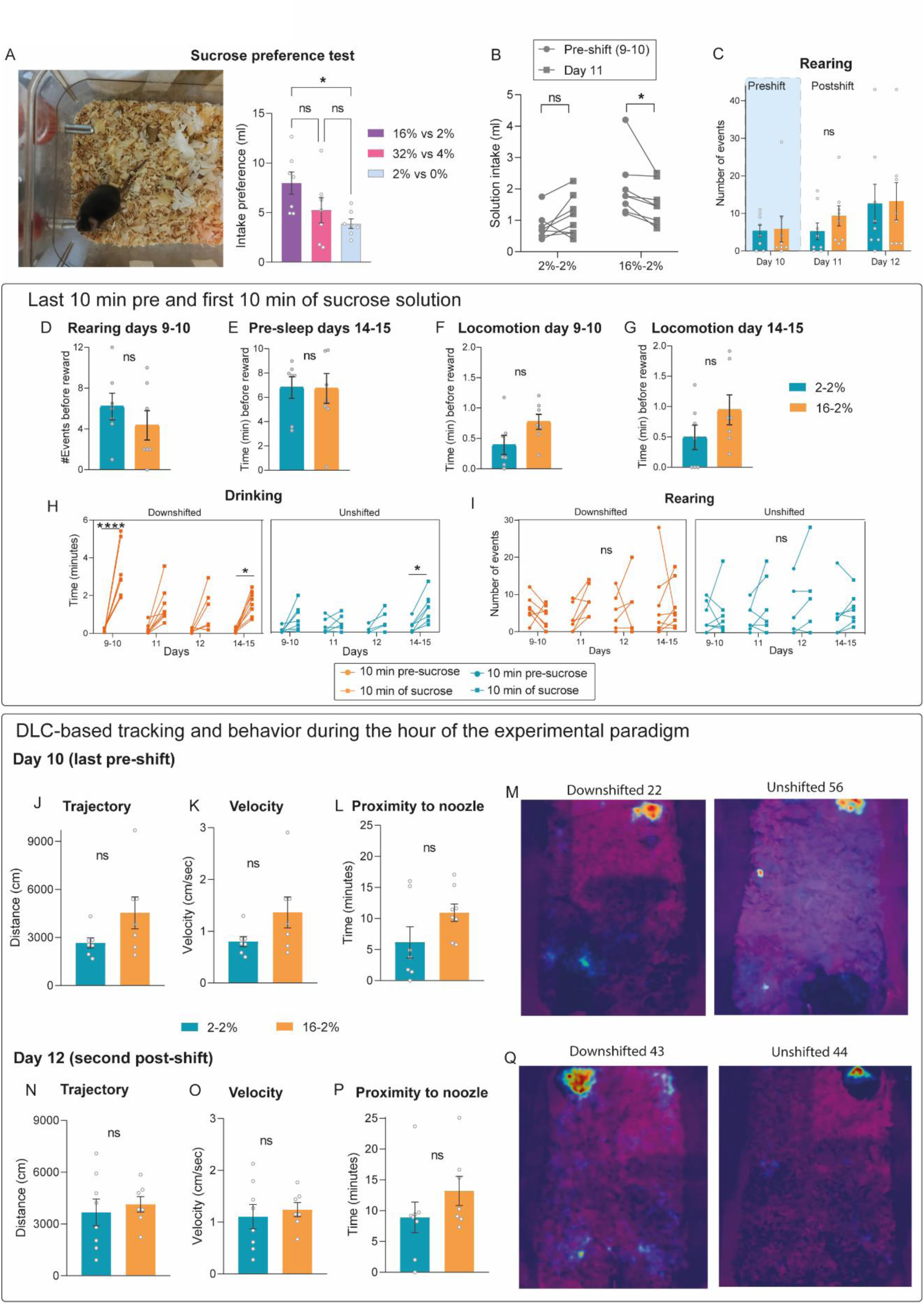
A) Photograph of the home-cage (left) used and results from the sucrose preference test (right), n=7, one-way RM ANOVA, p(Treatment) = 0.0481. See that two bottles are simultaneously presented in the cage and the animal can choose the preferred solution. B) Solution intake during the hour of the protocol on days 9-10 (mean value) and day 11 (first day post-shift) for downshifted (16-2%) and unshifted animals (2-2%), n=8, two-way RM ANOVA, p(Treatment) = 0.52, p(Treatment x Day) = 0.0067. C) Rearing behavior analyzed during the hour of the protocol for days 10-12, n=8 mice, two-way RM ANOVA, p(Treatment) = 0.64, p(Treatment x Day) = 0.7. D-G) Rearing, pre-sleep and locomotor behavior measured on days 9-10 and 14-15 during the 10 min pre-sucrose presentation. D) Rearing behavior on days 9-10, n=7 mice, unpaired t-test, p = 0.36. E) Pre-sleep on days 14-15, n=7 mice, unpaired t-test, p = 0.96. F) Locomotion on days 9-10, n=7 mice, unpaired t-test, p = 0.0787. G) Locomotion on days 14-15, n=7 mice, unpaired t-test, p = 0.18. H-I) Drinking and rearing behavior in the 10 min before reward and first 10 min of sucrose presentation. H) Drinking, n=7 mice, three-way RM ANOVA, p(Treatment) = 0.0078, p(Treatment x Day) = 0.0057, p(Treatment x 10min-window) = 0.0033, p(Treatment x x Day x 10min-window) = 0.0264. I) Rearing, n=7 mice, three-way RM ANOVA, p(Treatment) = 0.8, p(Treatment x Day) = 0.77, p(Treatment x 10min-window) = 0.86, p(Treatment x x Day x 10min-window) = 0.76. J-L) DLC-based tracking showing trajectory, velocity and time spent in the rewarded zone for downshifted and unshifted animals on day 10 (last pre-shift). J) Trajectory, n=7 mice, unpaired t-test, p = 0.097. K) Velocity, n=7 mice, unpaired t-test, p = 0.097. L) Proximity to nozzle, n=7 mice, unpaired t-test, p = 0.11. M) Heat maps of a downshifted (left) and an unshifted mouse (right) showing examples of their behavior during the hour of protocol on day 10. N-P) Trajectory, velocity and time spent in the rewarded zone on day 12 by downshifted and unshifted animals during the hour of the protocol. N) Trajectory, n=7 mice, unpaired t-test, p = 0.63. O) Velocity, n=7 mice, unpaired t-test, p = 0.63. P) Proximity to nozzle, n=7 mice, unpaired t-test, p = 0.24. Q) Heat maps of a downshifted (left) and an unshifted mouse (right) showing examples of their behavior during the hour of protocol on day 12. ns, p > 0.05; ∗, p < 0.05; ∗∗, p <0.01; ∗∗∗, p <0.001.

**Figure S2.**
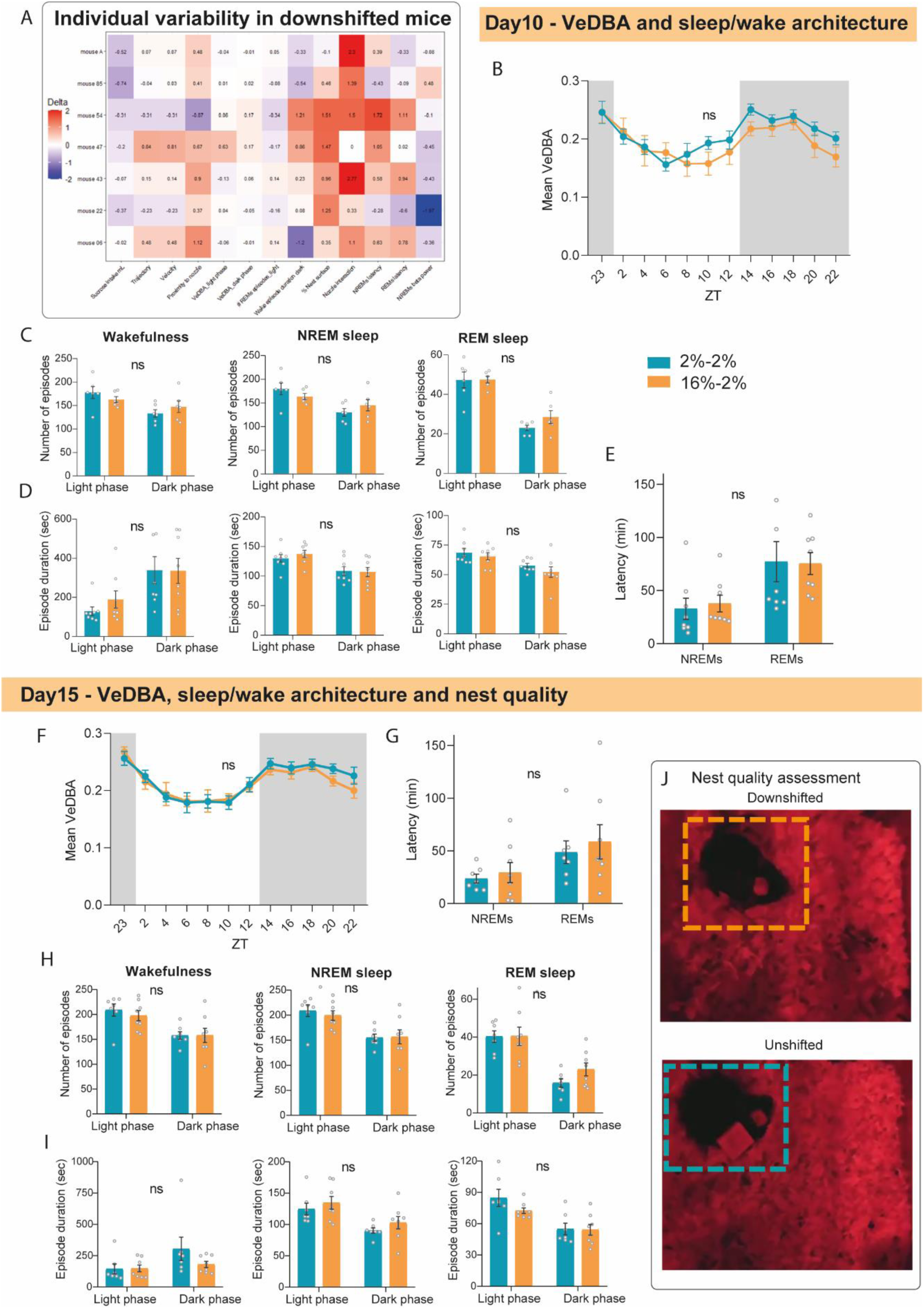
A) Individual differences for each of the downshifted mice (n=8) on day 11 relative to day 10 (values indicate log-ratio). B) VeDBA calculated for the hour of protocol and 2 hour means during day 10 (last pre-shift), n=8 mice, two-way RM ANOVA, p(Treatment) = 0.44, p(Treatment x Hour) = 0.49. C) Number of episodes of wakefulness, NREMs and REMs during the 12h of light phase and 10h of dark phase on day 10 (last pre-shift). Wakefulness, n=8 mice, two-way RM ANOVA, p(Treatment) = 0.8, p(Treatment x Phase) = 0.18. NREMs, n=8 mice, two-way RM ANOVA, p(Treatment) = 0.71, p(Treatment x Phase) = 0.13. REMs, n=8 mice, two-way RM ANOVA, p(Treatment) = 0.98, p(Treatment x Phase) = 0.29. D) Mean episode duration of wakefulness, NREMs and REMs during the 12h of light phase and 10h of dark phase on day 10. Wakefulness, n=8 mice, two-way RM ANOVA, p(Treatment) = 0.58, p(Treatment x Phase) = 0.58. NREMs, n=8 mice, two-way RM ANOVA, p(Treatment) = 0.7, p(Treatment x Phase) = 0.47. REMs, n=8 mice, two-way RM ANOVA, p(Treatment) = 0.2, p(Treatment x Phase) = 0.74. E) Latency to NREMs and REMs sleep from ZT0 and after the reward is removed from the cage on day 10, n=8 mice, two-way RM ANOVA, p(Treatment) = 0.92, p(Treatment x Stage) = 0.64. F-H) Same results as A-D) but for day 15 (last post-shift and end of experimental protocol). F) VeDBA calculated for the hour of protocol and 2 hour means during day 15 (last post-shift), n=8 mice, two-way RM ANOVA, p(Treatment) = 0.59, p(Treatment x Hour) = 0.82. F) Latency to NREMs and REMs sleep from ZT0 and after the reward is removed from the cage on day 15, n=8 mice, two-way RM ANOVA, p(Treatment) = 0.59, p(Treatment x Stage) = 0.77. G) Number of episodes of wakefulness, NREMs and REMs during the 12h of light phase and 10h of dark phase on day 15. Wakefulness, n=8 mice, two-way RM ANOVA, p(Treatment) = 0.68, p(Treatment x Phase) = 0.55. NREMs, n=8 mice, two-way RM ANOVA, p(Treatment) = 0.77, p(Treatment x Phase) = 0.56. REMs, n=8 mice, two-way RM ANOVA, p(Treatment) = 0.36, p(Treatment x Phase) = 0.31. I) Mean episode duration of wakefulness, NREMs and REMs during the 12h of light phase and 10h of dark phase on day 15. Wakefulness, n=8 mice, two-way RM ANOVA, p(Treatment) = 0.24, p(Treatment x Phase) = 0.25. NREMs, n=8 mice, two-way RM ANOVA, p(Treatment) = 0.34, p(Treatment x Phase) = 0.82. REMs, n=8 mice, two-way RM ANOVA, p(Treatment) = 0.22, p(Treatment x Phase) = 0.35. J) Example photographs used for nest quality assessment on day 15 for an unshifted and a downshifted mouse. ns, p > 0.05.

